# Disordered N-terminal region of CLK1 oligomerizes for recruitment to nuclear substructures and splicing function

**DOI:** 10.64898/2026.02.22.707342

**Authors:** Athira George, Laurent Fattet, Dominique Capraro, Patricia A. Jennings, Joseph A. Adams

## Abstract

Cdc2-like kinase 1 (CLK1) phosphorylates the serine-arginine (SR) proteins, a family of nuclear factors essential for mRNA splicing. The ability of CLK1 to recognize and efficiently modify SR proteins is strictly dependent on a lengthy, disordered N-terminus flanking its kinase domain. In addition to stimulating phosphorylation, this 150-residue extension also induces large oligomer formation in CLK1 but it is unclear whether said structure is important for catalytic or cellular function. We identified a subset of N-terminal residues that, upon removal, impairs oligomerization of CLK1 but does not abolish phosphorylation of the SR protein SRSF1. Despite robust phosphorylation, CLK1 lacking these high-order oligomerization sequences cannot effectively enter nuclear speckles and release SRSF1. This inability to mobilize CLK1 has detrimental effects on the alternative splicing of the CLK1 gene, severing an important, autoregulatory mechanism that controls active cellular levels of the kinase. Such findings indicate that whereas a limited group of residues in the N-terminus activates the kinase domain for SR protein phosphorylation, sequences that induce oligomerization direct CLK1 to the proper subnuclear structures for splicing function.

## Introduction

The Cdc2-like kinases (CLKs) phosphorylate and regulate the cellular properties of a critical group of splicing factors known as the serine-arginine (SR) proteins [1, 2]. This family of CLK substrates consists of twelve members (SRSF1-12) that contain one or two N-terminal RNA binding motifs (RRMs) and disordered, C-termini rich in Arg-Ser dipeptides known as RS domains [3, 4]. CLKs phosphorylate RS domains at both Arg-Ser and Ser-Pro dipeptides but the latter modifications are particularly important for controlling SR protein function in the nucleus. Ser-Pro phosphorylation has been shown to release the prototype SR protein SRSF1 (aka ASF/SF2) from membrane-less, nuclear compartments known as speckles so its RRMs may bind precursor mRNA (pre-mRNA) and help form the spliceosome, a macromolecular complex containing five small nuclear ribonuclear proteins (U1-6 snRNP) and numerous auxiliary proteins [5, 6]. SR proteins are essential for splicing based on *in vitro* complementation assays and have also been shown to play important roles in establishing splice sites in pre-mRNA [7–10]. All four human CLK isoforms (CLK1-4) possess conserved kinase domains (∼300 residues) and N-terminal extensions (∼150-300 residues) that are essential for high-affinity SR protein binding and Ser-Pro phosphorylation. Prior biophysical studies demonstrated that removal of the N-terminus from CLK1 significantly impairs SRSF1 binding affinity and Arg-Ser phosphorylation and completely abolishes critical Ser-Pro phosphorylation [11, 12]. The latter chemical transformation, readily identified by a decrease in SRSF1 migration on SDS-PAGE, generates a form commonly referred to as the hyper-phosphorylated or Hyper-P state. CLK1 expression has been shown to increase the Hyper-P state of SRSF1 in cells whereas CLK1 inhibition using ATP competitors such as TG003 has the opposite effect, lowering the Hyper-P state [13–15]. Owing to their ability to regulate splicing, CLK-directed inhibition is now being considered for the treatment of numerous diseases including cancer, Alzheimer’s disease, influenza A, diabetes and autism spectrum disorders [16–18].

The mechanism whereby CLKs target and regulate nuclear SR proteins has been evaluated using both *in vitro* biochemical and cell-based strategies. In addition to supporting high-affinity SR protein binding, the 150-residue N-terminus plays an important role in shifting CLK1 from the cytoplasm to the nucleus for SR protein phosphorylation using a “piggyback” mechanism. Lacking a classical nuclear localization sequence, we showed recently that CLK1 enters the nucleus by forming a complex with SRSF1 that binds to the transportin TRN-SR2 [19]. Formation of the SRSF1:TRN-SR2 complex for CLK1 nuclear transport, relies upon the cytoplasmic kinase SRPK1 that phosphorylates approximately eight Arg-Ser dipeptide repeats in the RS domain of SRSF1 providing the critical recognition element for TRN-SR2 [20–22]. In this manner, SRPK1 acts as a nuclear transport catalyst for both SRSF1 and CLK1. In the nucleus, SRPK1-phosphorylated SRSF1 constitutes a hypo-phosphorylated form (Hypo-P) that largely resides in speckles. Nuclear CLK1 can phosphorylate up to three Ser-Pro dipeptides in the RS domain, thereby generating the Hyper-P state, a transformation that dissociates SRSF1 from speckles for engagement with the developing spliceosome [6]. Thus, CLK1-dependent generation of Hyper-P induces conformational changes that release the phospho-RS domain from the neighboring RRMs in SRSF1 for pre-mRNA attachment [23]. Although CLK1 modifies SRSF1 for both speckle release and pre-mRNA binding, the intrinsic catalytic activity of this kinase is weak compared to other serine kinases such as SRPK1 [15, 24]. Furthermore, once phosphorylated, SRSF1 remains tightly bound to the N-terminus of CLK1 inhibiting splicing [11, 25]. Additionally, it has been shown that nuclear SRPK1 forms a complex with CLK1 enhancing both Ser-Pro phosphorylation and phospho-SRSF1 release from CLK1 [25]. The latter process involves severing high-affinity contacts between the RS domain of SRSF1 and the N-terminus of CLK1 using conserved docking residues on SRPK1 as an exchange platform [26]. Together, such findings demonstrate the importance of the N-terminus for CLK1 nuclear entry and engagement with the RS domains of SR proteins for phosphorylation-dependent splicing.

The CLK1 N-terminus not only recognizes SR proteins for catalysis but also regulates the quaternary structure of the kinase. While CLK1is a 57 kDa protein, it forms a large oligomer in solution with a hydrodynamic radius up to 80 nm and an apparent molecular mass of 1000 kDa or more [27]. In contrast, a form of CLK1 lacking its N-terminus (CLK1(ΔN)) has a hydrodynamic radius (4 nm) and apparent molecular mass (39 kDa) consistent with a monomer [27, 28]. This observed oligomerization phenomenon does not appear to be a result of the baculovirus expression system used to express and purify the recombinant enzyme since self-association was also observed between an RFP-tagged CLK1 and endogenous CLK1 in HeLa cells and between GST-and Myc-tagged CLKs in Cos cells [27, 29]. Unlike the kinase domain whose structure has been elucidated by X-ray diffraction methods [28], the CLK1 N-terminus is structurally disordered and is rich in both positively and negatively charged amino acids that could readily adapt to different surface charges on its target [25] (Fig1A). Indeed, CLK1 binds with high affinity to both the unphosphorylated and phosphorylated RS domain in SRSF1, in contrast to other protein kinases such as PKA and SRPK1 that readily dissociate their phospho-products owing to charge clashes in the active site [30, 31]. These observations raise the question of how a disordered, non-conserved N-terminus imparts changes in both CLK1 structure and hyper-P states of its SR protein substrate.

In this new study, we wish to understand how the N-terminus induces oligomerization of CLK1 and determine whether this unique quaternary structure is necessary for the catalytic and cellular function of the kinase. The N-termini within the CLK family are replete with charged and disorder-promoting residues and possess very low sequence conservation (<10%) (Fig1A). To investigate the function of the non-conserved N-terminus, deletions in CLK1 were generated with the larger goal of separating the oligomerization and catalytic properties of this domain. Using these deletion constructs, we found that a short stretch of residues (∼50 aa) directly flanking the kinase domain is sufficient for inducing Ser-Pro phosphorylation activity but does not induce large, oligomeric forms of CLK1. We discovered that a deletion form lacking oligomerization sequences catalyzes Ser-Pro phosphorylation *in vitro* but cannot efficiently enter into or release SRSF1 from nuclear speckles. The latter subcellular restraint has significant effects with regard to CLK1-dependent splicing activities. We found that whereas CLK1 expression induces exon 4 exclusion in its mRNA, a negative feedback mechanism that controls total CLK1 protein expression in cells [13, 32, 33], removal of the oligomerization sequences breaks this auto-regulatory loop. Such findings indicate that whereas induction of large oligomers through the N-terminus is not necessary for attaining the Hyper-P state of SRSF1, it is vital for controlling the subnuclear trafficking of CLK1 and its SR protein substrates. These results suggest that unlike other kinases with discreet domain regulation of activity (refs), CLK1 functionally partitions residues in its lengthy, disordered N-terminus for the initial purpose of localizing the enzyme near its substrate in the cell followed by positioning the RS domain of the SR protein for hyper-phosphorylation at discrete Ser-Pro dipeptides.

## Material and methods

### Materials

Tris-HCl, MOPS, NaCl, MgCl_2_, Ni-resin, GSH-resin, glutathione, DTT, IPTG, CHAPS, PMSF, triton X-100, Cellfectin II, liquid scintillant, imidazole, glycerol, Phenix imaging film and BSA were obtained from Fischer Scientific. FuGENE transfection reagent was obtained from Promega. Protease inhibitor cocktail was obtained from Roche. (γ-^32^P)-ATP was purchased from NEN Products. InstantBlue was purchased from Expedeon. Anti-His antibody resins were obtained from Santa Cruz Biotechnology. Anti-GST antibodies were obtained from BioLegend. All oligonucleotides were purchased from Integrated DNA Technologies. Anti-RFP and anti-GFP antibodies were obtained from Abcam. Anti-SC35 antibodies were obtained from Cell Signaling. Anti-actin antibodies were obtained from Sigma.

### Expression and purification of recombinant proteins

CLK1(ΔN) (UniProt Entry P49759), cloned into a pET28a vector, was purchased from Genscript and expressed in *E. coli* BL21(DE3) for 4 hours at 25°C after inducing with 0.5 mM IPTG. The cell pellet was re-suspended in lysis buffer (50 mM Tris HCl pH 7.5, 500 mM NaCl, 1 mM DTT and 0.04% CHAPS) and lysed by sonication at 25% power for 5 min with 15 s on and off cycles in the presence of protease inhibitor cocktail and PMSF. The lysate was incubated with pre-washed Ni-resin for 1 h at 4°C and then washed with lysis buffer (100 mL each) supplemented with 0, 5, and 30 mM imidazole. The protein was eluted using 300 mM imidazole, dialyzed overnight into 50 mM Tris-HCl (pH 7.5), 500 mM NaCl, 1 mM DTT, and 10% glycerol and stored in −80°C in small aliquots. CLK1, CLK1(Δ1), and CLK1(Δ12), cloned into pFastBacHTC vectors, were purchased from Genscript and transformed into DH10 Bac cells. Colonies expressing bacmids with CLK1 genes incorporated were selected with blue/white screening. The bacmids were purified from an overnight culture of the selected colony using purelink hipure plasmid miniprep kit and transfected into Sf9 cells plated in a 6-well plate at a density of 0.8 x 10^6^ cells/well. Cellfectin: DNA (4 μL cellfectin /1 μg DNA) complexes were added to Sf9 cells in serum free media. The baculoviruses were harvested 5 days after transfection and amplified before infecting Hi5 cells (density 2 x 10^6^ cells/mL) for protein expression. Cells were lysed and His-tagged proteins were purified using the same protocol as CLK1(ΔN). All GST-N constructs were cloned into pGEX vectors and expressed in BL21(DE3) *E. coli* using a 4 h induction at 25°C with 0.5 mM IPTG. The cells were lysed in 50 mM Tris-HCl (pH 7.5), 500 mM NaCl, 1 mM DTT and 0.025% triton X-100 in the presence of protease inhibitor cocktail and cells were spun down at 13,000 rpm at 4°C for 30 min. The supernatant was incubated with GSH-resin for one hour at 4°C. The resin was washed with 500 mL of lysis buffer before eluting with elution buffer containing 25 mM glutathione. The purified proteins were dialyzed against glutathione-free dialysis buffer (50 mM Tris-HCl (pH 7.5), 500 mM NaCl, 1 mM DTT, and 10% glycerol), analyzed using SDS-PAGE and stored at −80°C. GFP-SRSF1 and CLK constructs were expressed from pcDNA3.1+N-eGFP and pcDNA3-mRFP vectors in HeLa cells.

### Dynamic light scattering (DLS) and size exclusion chromatography

All DLS measurements were performed using a Protein Solutions DynaPro instrument and data were analyzed using the dynamics software. All samples were dialyzed into a buffer containing 50 mM Tris-HCl (pH 7.5), 500 mM NaCl, 1 mM DTT, 10% glycerol and spun down at 13,000 rpm for 10 min to remove any insoluble particles prior to data collection. The supernatant was transferred to a cuvette without introducing air bubbles. For each sample, 20 acquisitions were made after equilibrating the sample at 25°C. The R_H_ values were calculated using regularization analysis. Size exclusion chromatography was performed using an S200 column on a Biorad Duoflow FPLC system maintained at 4°C. The samples were run for 30 min at a flow rate of 0.7 mL/min in a running buffer containing 50 mM Tris-HCl (pH 7.5), 500 mM NaCl, and 1 mM DTT. The proteins were tracked by monitoring the absorbance at 280 nm. Molecular mass standards for the S200 columns were run under the same conditions as the samples.

### Phosphorylation kinetics studies

All single-turnover kinetic assays were performed in a buffer containing 100 mM MOPS, 22 mM Tris-HCl, 225 mM NaCl, 5 mg/mL BSA, 0.45 mM DTT, 4.5 % glycerol, and 10 mM MgCl_2_ at 37°C and pH 7.2. Reactions contained 100 μM ^32^P-ATP (4000-8000 cpm/mol) and 0.1 μM SRSF1 and were initiated with the addition of 1 μM enzyme. Reaction aliquots were withdrawn at various times, quenched in SDS-PAGE loading buffer and run on a 10% SDS-PAGE. The ^32^P-SRSF1 bands were cut from the dried gels and counts were quantified in the ^32^P channel on a scintillation counter. The reaction progress curves were fitted to single exponential functions. All steady-state kinetic assays were performed at 25°C in 100 mM MOPS (pH 7.2), 5 mg/mL BSA, 10 mM MgCl_2_, and 25 μM ATP. The concentrations of CLK1, CLK1(Δ1), CLK1(Δ12) were fixed at 100 nM and CLK1(ΔN) was fixed at 250 nM. The SRSF1 concentrations were varied from 50 to 2000 nM. The reactions were initiated by adding SRSF1 and then quenched with SDS-PAGE loading buffer after 5 min. The quenched samples were run on a 12% SDS-PAGE, dried overnight, and the ^32^P-SRSF1 bands were cut out and counted. The K_m_ and k_cat_ values were obtained by fitting the data to the quadratic function given in equation (1).

### Immunofluorescence and confocal imaging experiments

For imaging experiments, cells were washed with 1× PBS, fixed with 4% paraformaldehyde (PFA) for 20 minutes at room temperature, permeabilized with PBS-0.5% Triton X-100 for 10 minutes at 4°C, and then blocked with 20% goat serum in PBS (blocking buffer). Samples were incubated with primary antibody (overnight in blocking buffer), washed 3 times with 1× PBS, incubated with secondary antibodies for 1 hour at room temperature, washed 3 times with 1× PBS, and mounted in DAPI-containing mounting medium (Vector Laboratories). Confocal images were acquired using an Olympus FV1000 microscope. Images were linearly analyzed and pseudo-colored using ImageJ analysis software.

### Immunoprecipitation & pull-down experiments

For immunoprecipitation experiments, HeLa cell lysates were generated by incubating cells in 1×Ripa buffer at 4°C followed by sonication with protease inhibitor. Lysates (400 μl at 1 mg/ml), protease inhibitor, and immobilized antibody-bead conjugate (10 μl) were then incubated overnight with gentle rocking at 4 °C. The mixture was centrifuged at 1000 × *g* and washed 2× with 500 μl 1× PBS at 4 °C. The protein bands were visualized by Western immunoblotting. For pull-down experiments, GST-SRSF1 (1 μM) was incubated with His-tagged CLK1 proteins (10 μM) in binding buffer (0.1% NP40 (Nonidet P40), 20 mM Tris/HCl (pH 7.5) and 75 mM NaCl) in a total volume of 40 μL for 30 min before incubating with 25 μl of GSH-resin for 30 min at room temperature. In all cases, the resin was washed 3× with 200 μl of binding buffer and the bound proteins were eluted with SDS quench buffer and boiled for 5 min. Retained protein was resolved by 12% SDS-PAGE and visualized by InstantBlue Coomassie stain.

### Splicing assays

In order to monitor alternative splicing in HeLa cells expressing CLK1-RFP proteins, RNA was isolated from cell lysates using the RNeasy Plus kit from Qiagen. cDNAs were generated from the isolated RNA using the Superscript III Reverse transcriptase from Invitrogen. The alternatively spliced gene was then amplified using the GoTaq Green Master Mix from Promega using complementary oligonucleotides for sequences in exons 2 and 5. The amplified fragments were resolved on a 2% agarose gel and imaged using a Bio-Rad gel doc system. The following forward/reverse oligonucleotide set was used: CLK1 forward (exon 2): ATGAGACACTCAAAGAGAACTTAC; CLK1 reverse (exon 5): GCTTTATGATCGATGCACTCC.

## Results

### Isolating N-terminal sequences important for CLK1 oligomerization

Prior studies reveal that deletion of the full N-terminus shrinks CLK1 from a large oligomer to a small monomer [27] although it is unclear whether all of these residues or possibly a smaller subset are necessary for high-order oligomer formation. Since the N-terminus is a highly disordered domain with few conserved residues and numerous charged side chains similar to RS domains (Fig 1A), we elected to explore its function in a region-specific manner. The N-terminus of CLK1was divided into three separate blocks of approximately 50-residues each (Fig 1B). We expressed CLK1, CLK1(Δ1), and CLK1(Δ12) in insect cells and the kinase domain lacking the N-terminus, CLK1(ΔN), in *E. coli* and purified them using Ni-affinity chromatography (Fig 1C). All four proteins migrated at their calculated molecular masses on SDS-PAGE (61, 54, 48, and 42 kDa for CLK1, CLK1(Δ1), and CLK1(Δ12) and CLK1(ΔN), respectively). To assess their structural characteristics, we initially performed dynamic light scattering (DLS) and found that CLK1 displayed a hydrodynamic radius (R_H_) of 42 nm, consistent with a large oligomer, while the R_H_ for CLK1(ΔN) was only 4.8 nm (Fig1 D, E). These findings are consistent with previously published results showing that the N-terminus induces transition from a monomeric to a large oligomeric form of CLK1 [27]. Although removal of block 1 (residues 1-50) in CLK1(Δ1) modestly reduces R_H_ by only 33% compared to wild-type (28 vs. 42 nm), removal of blocks 1 and 2 (residues 1-104) in CLK1(Δ12) led to a significant decrease in R_H_ of almost 7-fold compared to wild type (6.2 vs 42 nm) (Fig1D, E). Indeed, the R_H_ for CLK1(Δ12) is very similar to that for CLK1(ΔN) (Fig1E). These findings suggest that residues in block 3 are insufficient for inducing high-order oligomer formation in CLK1.

**Fig 1.**
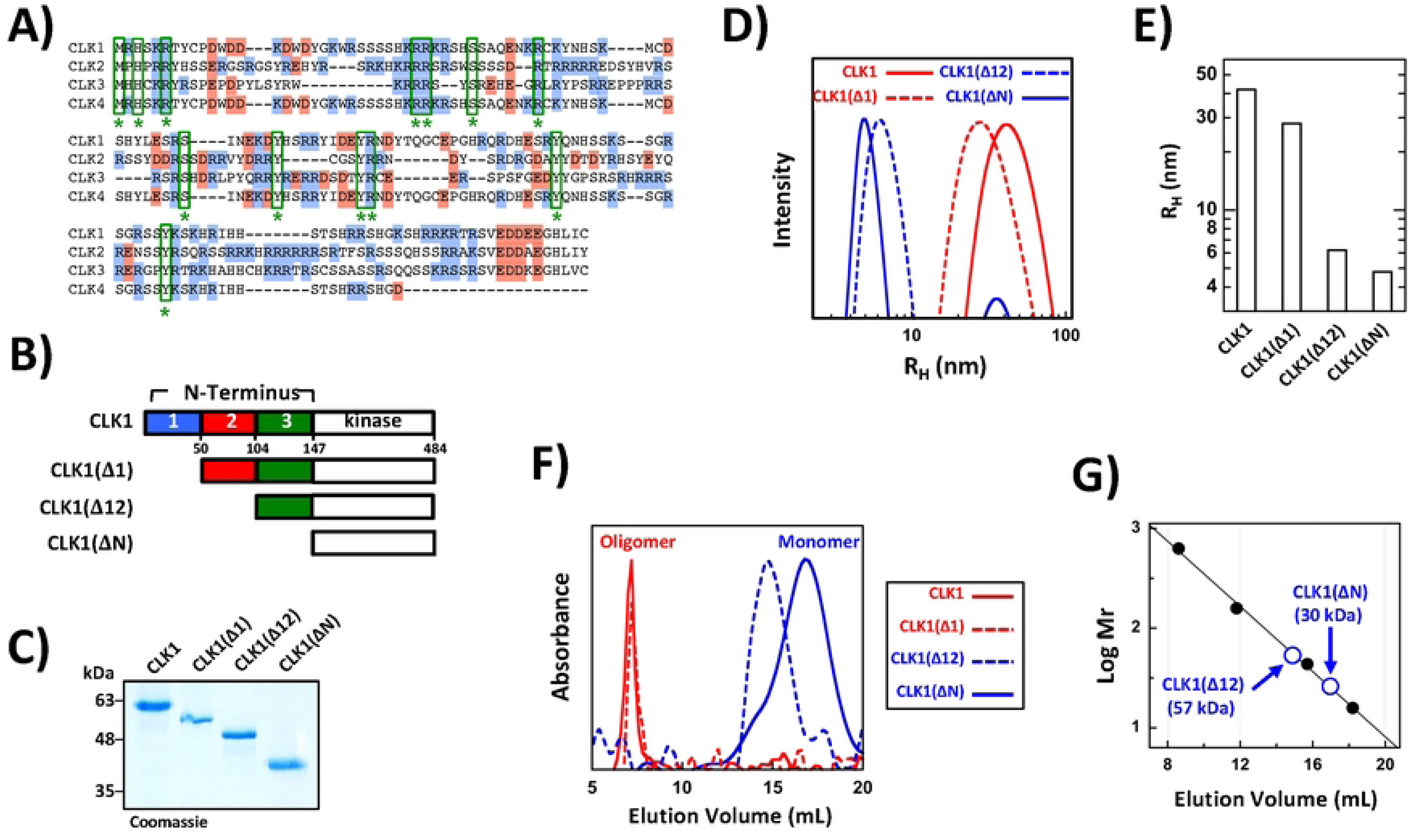
N-terminal regions controlling CLK1 oligomerization. A) Overlay of the N-termini from human CLK1-4. Sequences 1-151, 1-153, 149-294, and 1-131 in CLK1, CLK2, CLK3, and CLK4, respectively, are aligned. Conserved residues are denoted with asterisks and enclosed in green boxes and positively and negatively charged side chains are highlighted in blue and red. B) CLK1 deletion constructs. C) SDS-PAGE of CLK1 and the deletion constructs purified by Ni-resin affinity chromatography. D) DLS of CLK1 and deletions constructs. Total protein concentrations are 7 μM for CLK1 and CLK1(Δ1) and 30 μM for CLK1(Δ12) and CLK1(ΔN). E) Hydrodynamic radii (R_H_) of CLK1 and deletion constructs obtained from DLS measurements. F) SEC of CLK1 and deletions constructs run on an S200 column. G) SEC standardization curve. Molecular masses (Mr) of standard proteins (black, filled circles) are plotted against their elution volumes. The elution volumes and corresponding Mr values for CLK1(ΔN) and CLK1(Δ12) are displayed.

To further investigate the role of the N-terminus, we next performed size exclusion chromatography (SEC). We found that CLK1 and CLK1(Δ1) eluted from an S200 column just prior to the highest molecular weight standard implying that both proteins exhibit apparent molecular masses of, at least, 600 kDa, consistent with the large, hydrodynamic radii obtained from the DLS experiments (Fig1 F, G). As expected, CLK1(ΔN) displayed an apparent molecular mass (30 kDa) consistent with a monomer. In comparison, CLK1(Δ12) eluted with an apparent molecular mass (57 kDa) that is approximately twice that observed for CLK1(ΔN), a result consistent with a dimer or possibly an elongated monomer (Fig1 F, G). An additional assessment of the major peaks from SEC by SDS-PAGE confirmed the CLK1 constructs proteins were the correct molecular masses rather than breakdown products (SI Fig 1A). The combined data from separate DLS and SEC experiments indicate that residues in block 3 in the N-terminus do not support large particle formation but only induce dimers or possibly elongated monomers. Instead, addition of block 2 appears to play a significant role in inducing high-order oligomerization of CLK1 with block 1 addition providing further, albeit less significant, enhancement.

### Oligomerization region in the CLK1 N-terminus

Our structural studies suggest that block 3 in the N-terminus may induce a possible monomer-to-dimer transition whereas the further addition of blocks 1 & 2 induces high-order oligomerization of CLK1. To better understand how these blocks may seed this quaternary structural change, we studied the N-terminus of CLK1 in the absence of the kinase domain. We expressed and purified the N-terminus and several block deletions complementing the CLK1 deletions as GST fusion proteins in *E. coli* (Fig2 A, B). All constructs migrated near their calculated molecular masses based on SDS-PAGE (44 kDa for GST-N, 39 kDa for GST-N(Δ3), and 34 kDa for GST-N(Δ23) and GST-N(Δ13)) indicating that the N-termini constructs are not cleaved from the GST carrier protein. Unfortunately, we were not able to purify GST constructs containing only block 3 or block 3 attached to block 2 for comparison.

Using DLS, we assessed the hydrodynamic radius, R_H_, of the individual constructs, to determine the differences in size. Generally, the DLS experiments revealed that GST-N has an R_H_ of about 70 nm, consistent with a large oligomer (Fig2C,D). Interestingly, removal of block 3 in GST-N(Δ3) also resulted in a large R_H_ value (30 nm) suggesting that these final residues are not essential and the block 1-2 combination is sufficient for high-order oligomerization of the N-terminus (Fig2 C, D). In comparison, both single-block constructs (GST-N(Δ13) and GST-N(Δ23)) possess R_H_ values close to that for free GST indicating that both blocks 1 and 2 are necessary for self-association in GST-N(Δ3) (Fig2 C, D). For GST-N(Δ13), we detected a small peak with an R_H_ approximate to that for GST-N. We suspect that this peak constitutes a minor component of the total protein sample since DLS is highly sensitive in detecting large, oligomeric compared to small protein species. This suggests that whereas block 1 plays a minor role in CLK1 oligomerization (Fig1), it is essential in context of a shorter N-terminal fragment that lacks block 3. Likewise, block 2, which induced high-order oligomerization when attached to block 3 and the kinase domain, did not form oligomers alone. These findings indicate that while the CLK1 N-terminus can self-associate (N-N contacts) to form large oligomers, a single block is not sufficient for such quaternary structure.

Having demonstrated that a minimal combination of blocks 1 and 2 is capable of engaging in N-N contacts based on DLS, we next explored these structural characteristics using SEC. We found that both GST-N and GST-N(Δ3) eluted as large oligomers (>600 kDa) on an S200 column suggesting that the CLK1 N-terminus can self-associate in the absence of block 3 (Fig2E). Interestingly, we also observed an additional peak for each protein sample that eluted slightly before free GST. This elution peak for GST-N(Δ3) contains the full-length protein and does not constitute a breakdown product but rather represents a reduced quaternary state of the protein (SI Fig 1B). These findings suggest that the N-terminus can engage in highly dynamic, N-N contacts. In order to evaluate whether disulfide bonds are driving oligomerization, we incubated GST-N(Δ3) in the presence of a strong reducing agent (50 mM TCEP) and found that it did not affect protein oligomerization compared to a sample lacking pre-treatment (SI Fig 1C). Such findings indicate that N-N contacts are not driven by cysteine cross-bridging in the N-terminus. Importantly, compared to GST-N(Δ3), the major peaks for both GST-N(Δ13) and GST-N(Δ23) eluted with molecular masses close to that of free GST and lacked a higher mass fraction indicating that neither individual block is capable of inducing oligomers, consistent with the DLS analyses (Fig2 E, F). As expected, free GST eluted as a dimer with an apparent molecular mass of about 50 kDa (Fig2F). These experiments reveal that the CLK1 N-terminus forms large oligomers in solution in the absence of the kinase domain and residues in block 3 are not necessary for oligomerization. Interestingly, while a combination of blocks 1 and 2 can induce oligomerization in the absence of the kinase domain, the combination of blocks 2 and 3 are sufficient to induce large oligomers in the presence of the kinase domain. Such findings suggest that individual blocks may function in a flexible manner to induce large-scale CLK1 oligomer formation but a minimum of two blocks is necessary for such quaternary structure.

### N-terminal sequences are necessary for Ser-Pro hyper-phosphorylation activity

Additionally, we explored how the N-terminus influences catalysis at a region-specific level by monitoring the phosphorylation kinetics of SRSF1 using the CLK1 deletions. In prior studies we showed that while CLK1 can phosphorylate Arg-Ser dipeptides similar to SRPK1, the specific targeting of three critical Ser-Pro dipeptides in the C-terminal portion of the RS domain directly regulates the subnuclear localization and splicing functions of SRSF1 [6] (SI Fig 2). In single-turnover kinetic experiments where the enzyme concentration exceeds that of SRSF1, we found that CLK1 induced a prominent gel shift in SRSF1 on an SDS-PAGE autoradiogram whereas CLK1(ΔN) did not (Fig3A). For CLK1, this result is due to phosphorylation of three Ser-Pro dipeptides in the RS domain of SRSF1 that induces slower migration on SDS-PAGE (Hyper-P), as previously found [12]. Furthermore, although the N-terminus is essential for attaining

**Fig 2.**
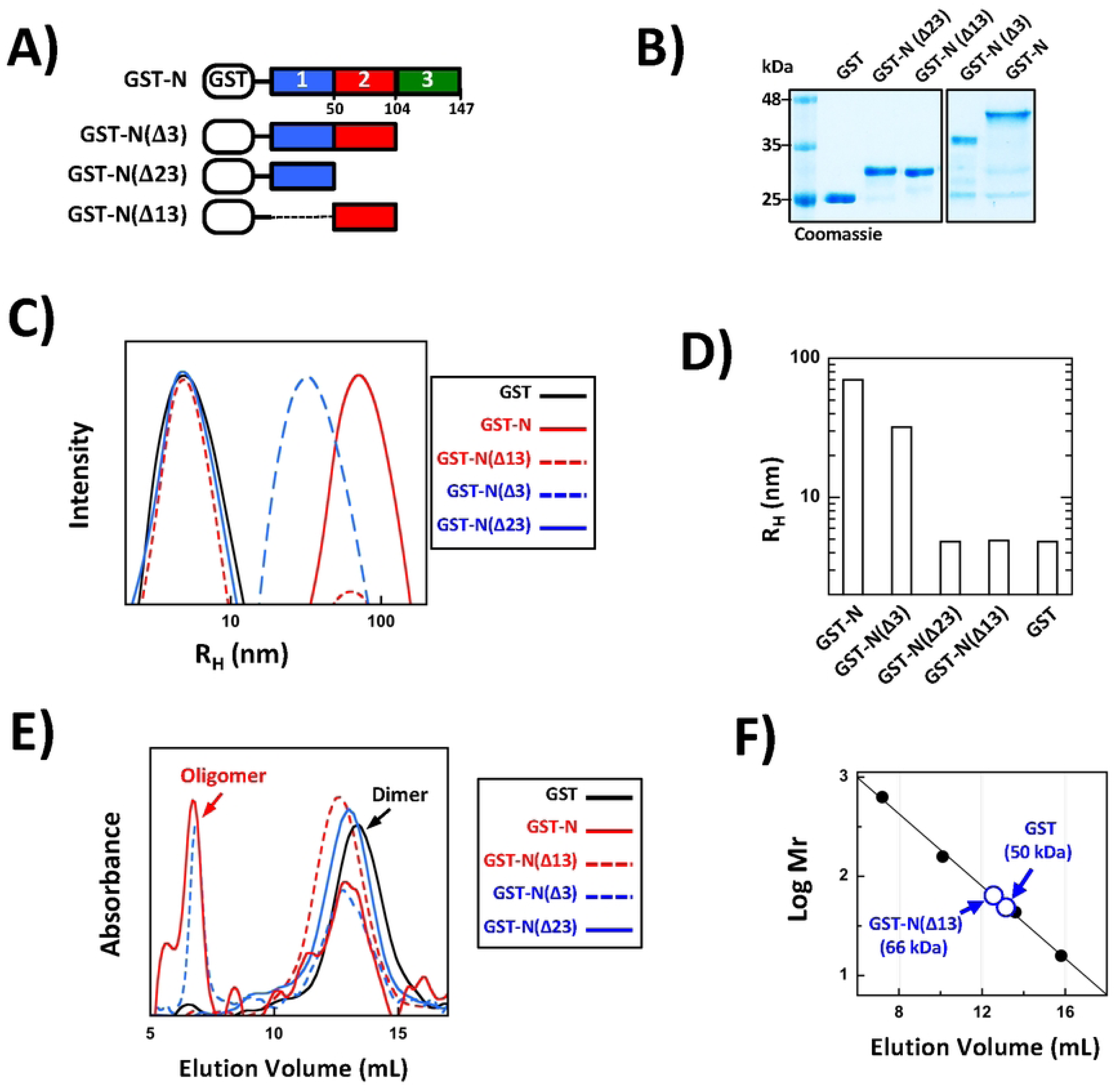
Deletion analyses of the CLK1 N-terminus. A) Deletion forms of the CLK1 N-terminus fused to GST. B) SDS-PAGE of GST-N and deletion constructs. C) DLS of GST-N and deletion constructs. Total protein concentrations are 3 μM for GST-N, 20 μM for GST-N(Δ3) and 50 μM for GST-N(Δ13), GST-N(Δ23) and GST. D) Hydrodynamic radii (R_H_) of GST-N and deletions obtained from DLS measurements. E) SEC of GST-N and deletions run on an S200 column. F) SEC standardization curve. Molecular masses (Mr) of standard proteins (black, filled circles) are plotted against their elution volumes. The elution volumes and corresponding Mr values for GST and GST-N(Δ13) are displayed.

**Fig 3.**
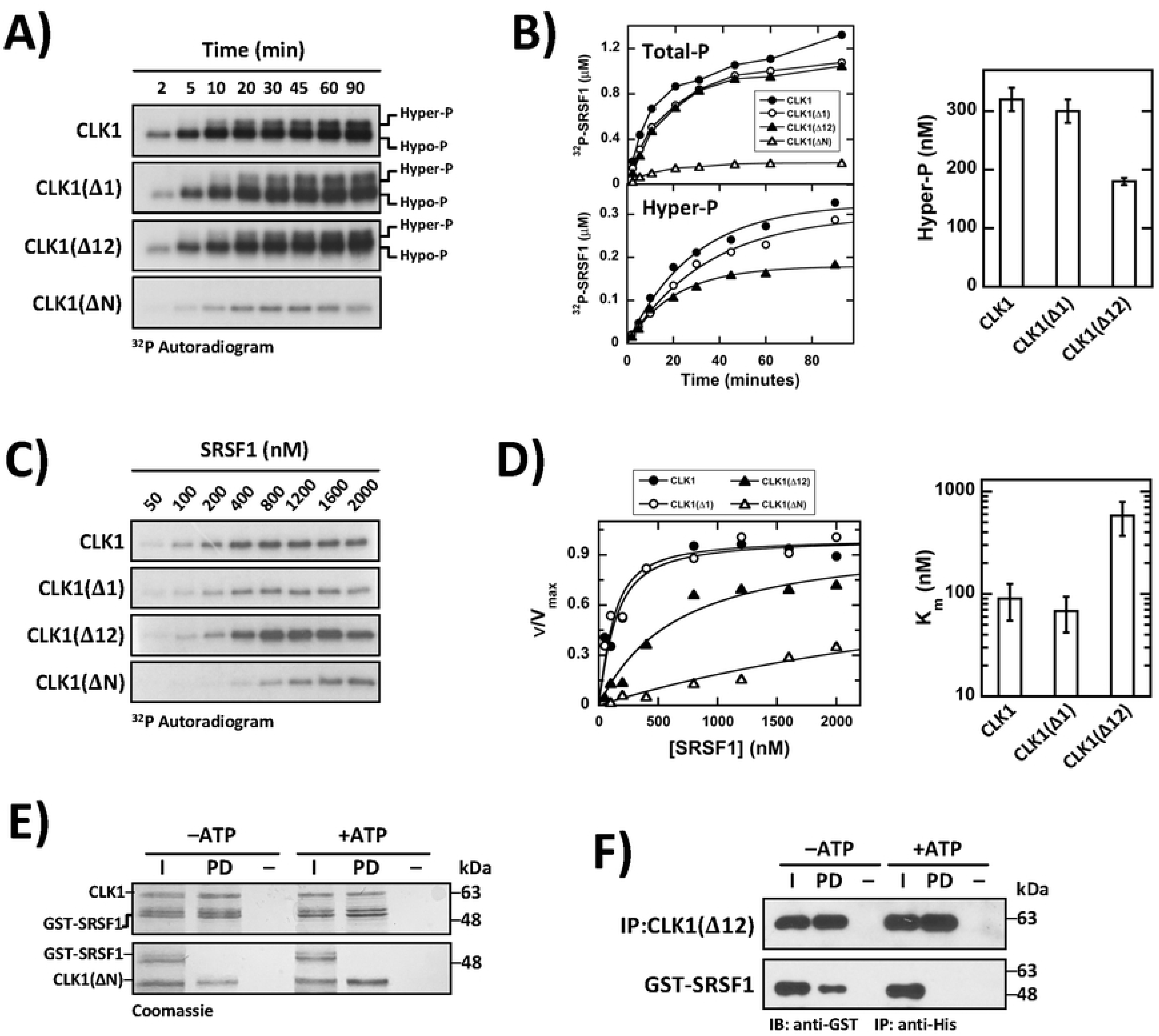
Enzyme kinetic analyses of CLK1 and deletion constructs. A) Single-turnover kinetic experiments. Autoradiograms for time-dependent SRSF1 phosphorylation by CLK1 and deletions forms are displayed. Enzymes (1 μM) are preincubated with SRSF1 (100 nM) and reacted with ^32^P-ATP (100 μM) for varying times. B) Total phosphorylation (Total-P) and hyper-phosphorylation (Hyper-P) from single-turnover kinetic experiments are plotted. Each time point is an average of two separate experiments (n=2). Maximum Hyper-P for each kinase form is plotted in the bar graph with error bars representing ±SD from data fitting. C) Steady-state kinetic experiments. Autoradiograms for SRSF1 phosphorylation are displayed using CLK1 and deletions forms. CLK1, CLK1(Δ1) and CLK1(Δ12) at 100 nM and CLK1(ΔN) at 250 nM are preincubated with varying SRSF1 and reacted with ^32^P-ATP (25 μM) for 5 minutes. D) Plots of initial velocity normalized to V_max_ (v/V_max_) vs total SRSF1 concentration. At each SRSF1 concentration, two separate experiments were averaged (n=2). Data are fit to equation (1) to obtain V_max_ values of 8.4 ± 0.55, 6.0 ± 0.34 and 21 ± 2.6 min^-1^ and K_m_ values of 90 ± 35, 68 ± 26, and 580 ± 210 nM for CLK1, CLK1(Δ1) and CLK1(Δ12), respectively. K_m_ values are plotted in the bar graph and error bars represent ±SD from data fitting. For CLK1(ΔN), kinetic data are normalized to a lower limit of 15 min^-1^ for V_max_. E) N-terminus of CLK1 is necessary for interactions with SRSF1. CLK1 and CLK1(ΔN), immobilized on Ni-agarose, are incubated with GST-SRSF1 in the absence and presence of ATP, washed and run on SDS-PAGE. The displayed gel is a representiative of two separate experiments with similar results (n=2). F) Oligomerization subdomain facilitates binding of phospho-SRSF1 to CLK1. CLK1(Δ12), immobilized on Ni-agarose resin, is incubated with GST-SRSF1 in the absence and presence of ATP, washed and run on SDS-PAGE. CLK1(Δ12) and GST-SRSF1 are detected using anti-His and anti-GST antibodies. The displayed gel is a representative of two separate experiments with similar results (n=2).

Hyper-P, it also strongly activates Arg-Ser phosphorylation since CLK1(ΔN) is deficient in overall phosphorylation compared to CLK1 (Fig3A,B). In the observed time frame of the single-turnover experiments, CLK1 added approximately 13 phosphates based on observed ^32^P incorporation and total SRSF1 concentration ([^32^P-SRSF1]/[SRSF1]_tot_) whereas CLK1(ΔN) added only 2 phosphates. Importantly, we found that removal of block 1 or blocks 1 and 2 did not abolish the generation of the Hyper-P state (Fig3B). These findings indicate that residues in block 3, while inadequate for inducing large CLK1 oligomers, are sufficient to activate the flanking kinase domain for SRSF1 hyper-phosphorylation. Such findings allow us to separate residues in the N-terminus that induce high-order oligomers of CLK1 from those that induce Ser-Pro phosphorylation.

### Oligomerization stabilizes SRSF1 binding for enhanced hyper-phosphorylation

Although block 3 in CLK1(Δ12) is sufficient for the induction of SRSF1 hyper-phosphorylation, the addition of block 2 in CLK1(Δ1) that supports high-order oligomerization enhances the extent of this critical CLK1-specific modification (Fig3B). We next wished to determine whether such enhancement is linked to increased stability of the CLK1:SRSF1 complex. To address this, we performed steady-state kinetic analyses using CLK1 and the deletion constructs. We mixed CLK1, CLK1(Δ1), and CLK1(Δ12) at 100 nM with varying SRSF1 (100-2000 nM) and allowed the reactions to incubate for 5 minutes prior to quenching (Fig3C). For CLK1(ΔN), we used higher enzyme concentrations (250 nM) owing to its lower, observed activity. At all SRSF1 concentrations, less than 6% of the substrate is phosphorylated based on observed ^32^P incorporation, thereby ensuring appropriate measurements of initial velocities in these experiments. However, since the enzyme levels needed to obtain sufficient ^32^P incorporation over the assay time frame were comparatively high, we fitted plots of initial velocity versus total substrate concentration using equation 1:

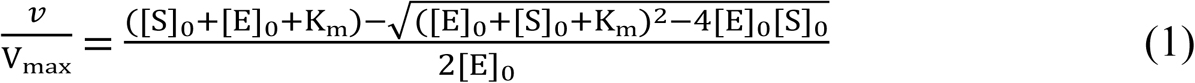

to obtain K_m_ values for SRSF1 [34] (Fig3D). In equation (1), [S]_o_ and [E]_o_ are the total SRSF1 and kinase concentrations and *v* and V_max_ are the initial and maximal velocities. We found that removal of the N-terminus increased the K_m_ more than 20-fold (100 to >2000 nM) indicating that this domain is not only vital for Ser-Pro hyper-phosphorylation but also for high-affinity binding of the SR protein. This observation is consistent with a prior report comparing steady-state kinetic profiles of CLK1 and CLK1(ΔN) [27]. Whereas removal of block 1 in CLK1(Δ1) had no observed effect, we found that removal of blocks 1 and 2 in CLK1(Δ12) raised the K_m_ by approximately 6-fold. These findings indicate that while residues directly flanking the kinase domain in block 3 allow the kinase domain to catalyze intrinsic hyper-phosphorylation, residues outside this block provide additional stability that correlate with increased Ser-Pro phosphorylation.

Although our steady-state kinetic analyses indicate that the addition of residues in block 2 of the N-terminus enhances SRSF1 affinity, these data report on apparent affinities (i.e.- K_m_ values) early in the reaction and prior to Ser-Pro dipeptide modification that is associated with the Hyper-P state. To evaluate SRSF1 affinities along the polyphosphorylation time course, we performed pull-down assays as a function of ATP phosphorylation. Similar to previous reports [25], we found that CLK1, immobilized on a Ni-agarose resin, interacted favorably with GST-SRSF1 either in the absence or presence of extensive ATP incubation suggesting that the substrate maintains high affinity during numerous phosphorylation events in the RS domain (Fig3E). As expected, CLK1(ΔN) did not interact favorably with GST-SRSF1 either in the absence or presence of ATP consistent with its poor binding affinity. We next explored whether CLK1(Δ12) also remains tightly bound to the substrate during phosphorylation. Since CLK1(Δ12) migrates close to GST-SRSF1 on SDS-PAGE we followed binding efficiency using Western blot analyses. We found that whereas CLK1(Δ12) interacted with GST-SRSF1 in the absence of ATP, it did not interact favorably after ATP incubation (Fig3F). These findings suggest that the affinity of GST-SRSF1 for CLK1(Δ12) decreases as a function of progressive phosphorylation. Thus, it is possible that the greater hyper-phosphorylation levels of CLK1 compared to CLK1(Δ12) may be due to enhanced stability of the SR protein on CLK1 not only at the onset of the reaction where mostly Arg-Ser dipeptides are engaged but also during the later stages where Ser-Pro dipeptides are modified for induction of the Hyper-P state.

### Residues controlling CLK1 subnuclear localization

Since we were able to abolish large oligomeric forms of CLK1 yet maintain critical SRSF1 hyper-phosphorylation by removing the first one hundred residues in the N-terminus (blocks 1 and 2), we next wished to investigate whether these sequences can regulate CLK1 subnuclear localization. To accomplish this, we generated CLK1, CLK1(Δ12) and CLK1(ΔN) constructs with C-terminal RFP tags for imaging in HeLa cells using confocal microscopy. These constructs allow us to evaluate multiple quaternary states, namely the large oligomeric (CLK1-RFP), monomeric/dimeric (CLK1(Δ12)-RFP) and monomeric (CLK1(ΔN)-RFP) forms of CLK1. We found that CLK1-RFP is expressed exclusively in the nucleus of HeLa cells whereas CLK1(Δ12)-RFP and CLK1(ΔN)-RFP are expressed in both the cytoplasm and nucleus, consistent with prior results indicating that the N-terminus facilitates nuclear localization [19, 29] (Fig 4A). In the nucleus, we found that CLK1-RFP displays greater localization in granular structures compared to the deletion constructs (Fig4B). These structures largely correspond with speckles based on staining for the speckle marker protein SC35 (Fig4A). Removing either the first 100 residues or the full N-terminus significantly reduced granular localization compared to wild-type CLK1 (Fig4B). Interestingly, either deletion that abolished the large oligomeric form of CLK1 (CLK1(Δ12)-RFP or CLK1(ΔN)-RFP) resulted in similar decreases in granular occupancy. These findings indicate that while the full N-terminus is important for efficient cytoplasmic-to-nuclear translocation, sequences in the N-terminus that induce large oligomers are critical for efficient CLK1 entry into nuclear speckles.

**Fig 4.**
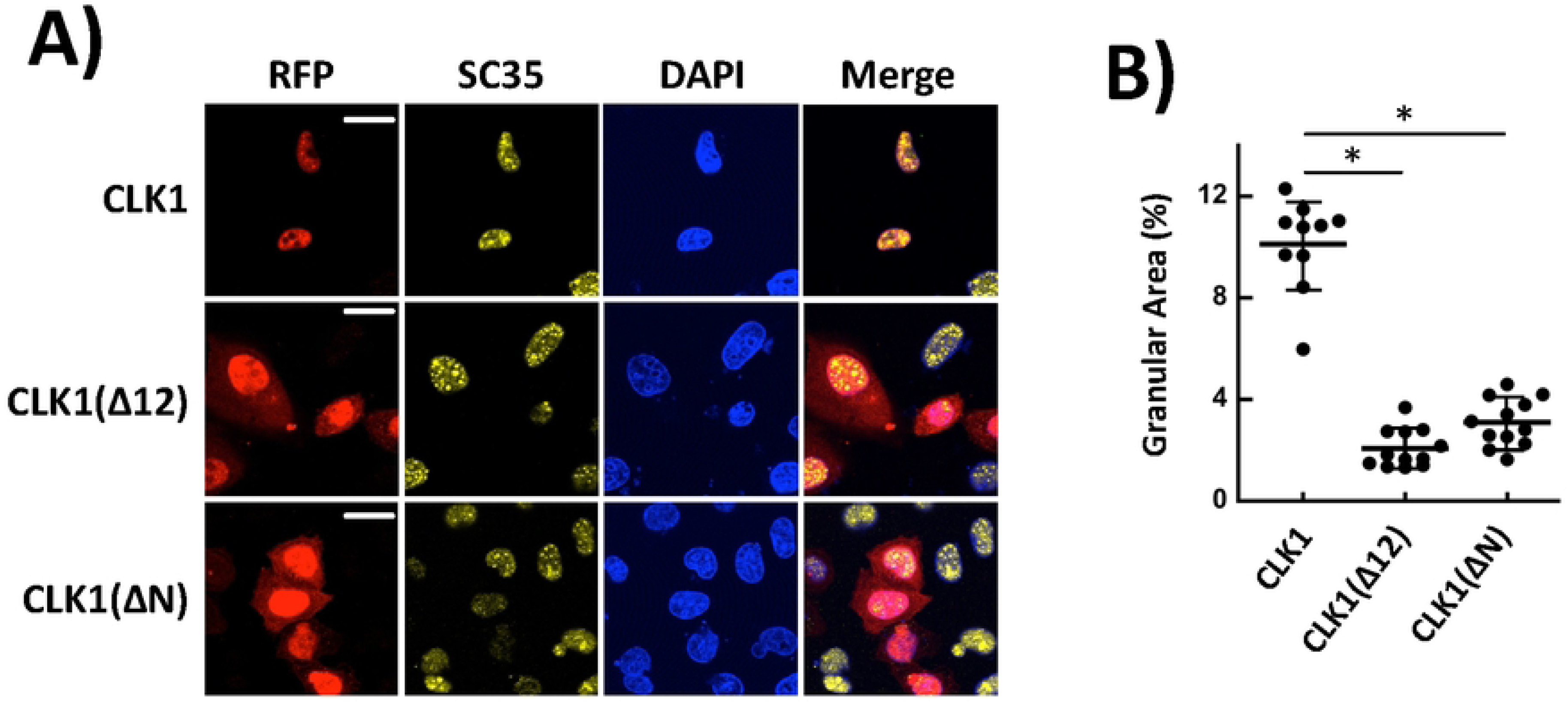
Oligomerization sequences control CLK1 subnuclear localization. A) Confocal microscopy of CLK1-RFP, CLK1(Δ12)-RFP and CLK1(ΔN)-RFP expressed in HeLa cells. RFP is monitored using direct fluorescence and SC35 is monitored using immunofluorescence. Scale bars, 20 μm. DAPI, 4,6-diamino-2-phenylindole. B) Granular nuclear areas for CLK1 constructs. Areas are calculated using IMAGEJ and displayed as dot plots with mean values ±SD where each dot represents analysis of an individual cell. P-values are calculated using a one-way ANOVA test (* is P<.0001).

**Fig 5.**
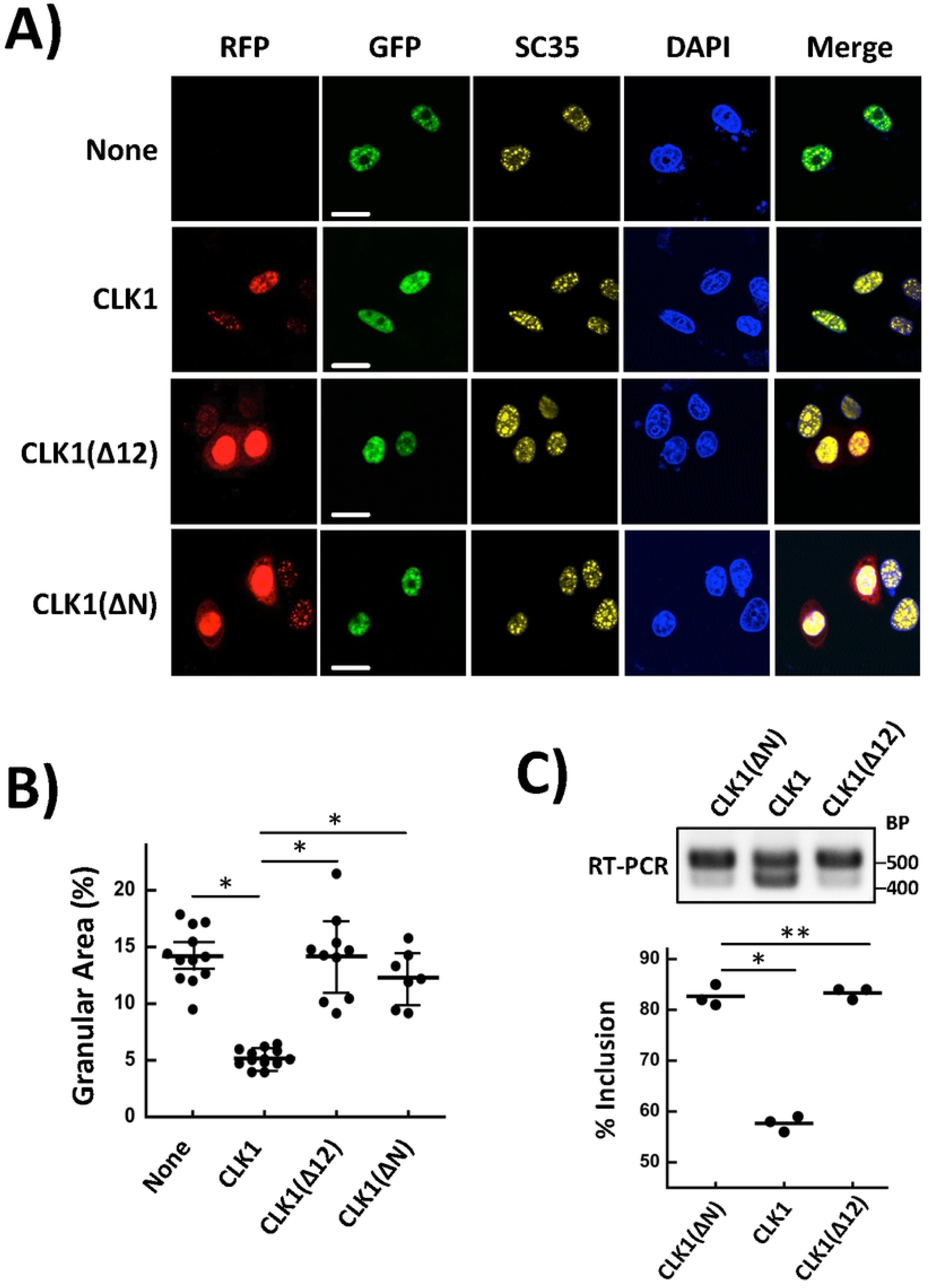
Effects of N-terminal deletions in CLK1 on SRSF1 subnuclear localization and splicing. A) Confocal microscopy of GFP-tagged SRSF1 expressed in HeLa cells as a function CLK1-RFP, CLK1(Δ12)-RFP and CLK1(ΔN)-RFP co-expression. GFP and RFP are monitored using direct fluorescence and SC35 is monitored using immunofluorescence. Scale bars, 20 μm. DAPI, 4,6-diamino-2-phenylindole. B) Granular nuclear areas for GFP-SRSF1. Areas are calculated using IMAGEJ and displayed as dot plots with mean values ±SD where each dot represents analysis of an individual cell. P-values are calculated using a one-way ANOVA test (* is P<.0001). C) Alternative splicing of exon 4 in the *CLK1* gene with CLK1 expression. RT-PCR is performed on RNA samples from HeLa cells expressing CLK1-RFP and deletion constructs.

**Fig 6.**
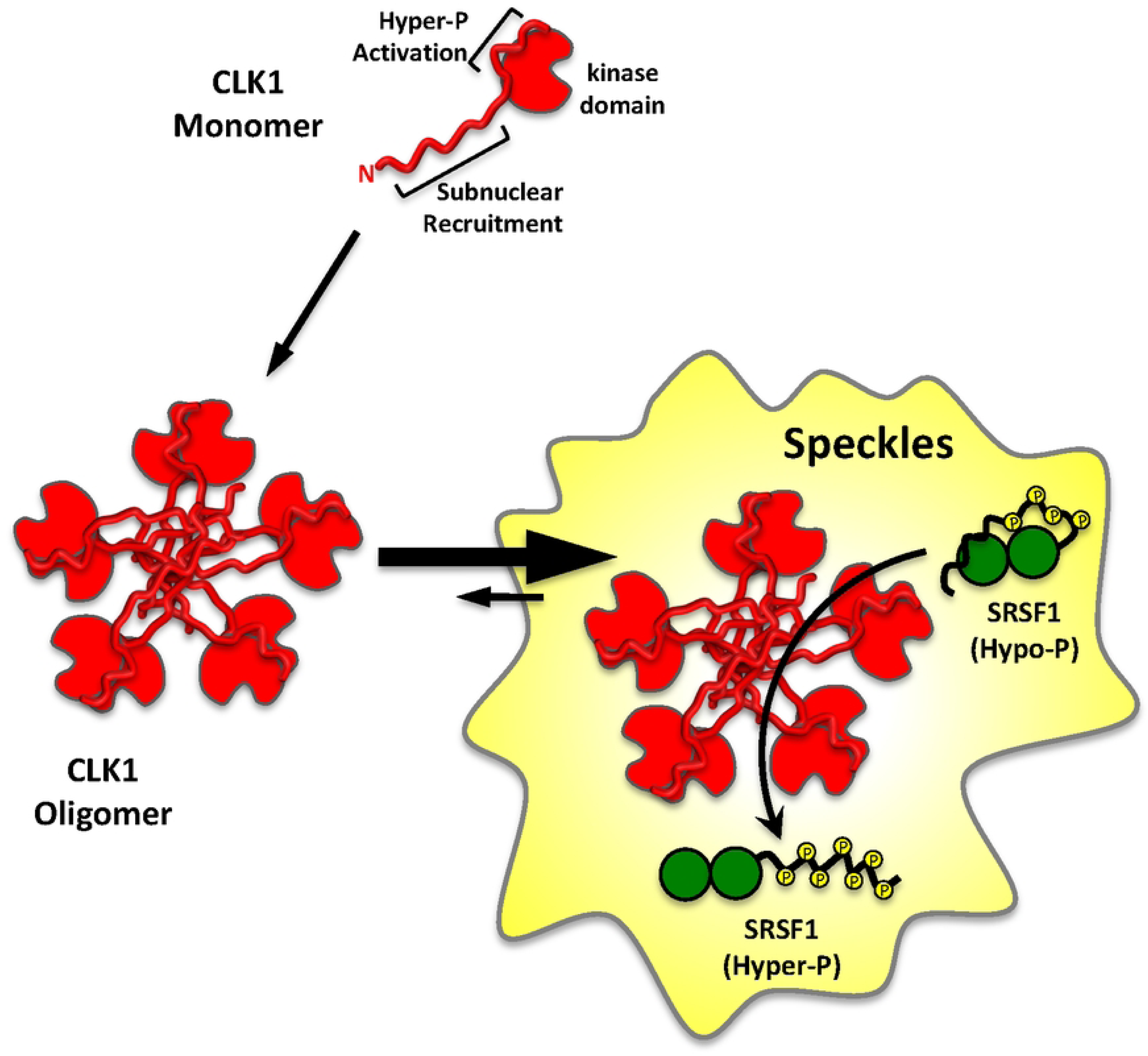
Model for CLK1 subnuclear localization and SR protein phosphorylation through N-terminal sequences. Subnuclear recruitment sequences in the N-terminus direct CLK1 to the nuclear speckles through high-order oligomer formation. Sequences directly flanking the kinase domain induce hyper-phosphorylation activity of the kinase domain.

### CLK1 oligomerization facilitates SRSF1 release from nuclear speckles

Since oligomerization sequences in the N-terminus support CLK1 entry into nuclear speckles (Fig4), we next considered whether such recruitment plays a regulatory function for SR proteins. To address this, we expressed GFP-SRSF1 in HeLa cells and monitored the effects of CLK1-RFP, CLK1(Δ12)-RFP and CLK1(ΔN)-RFP co-expression on the subnuclear localization of the SR protein (Fig5A). We found that CLK1-RFP expression greatly increased dispersion of GFP-SRSF1 from nuclear speckles to the nucleoplasm (14 vs 4.8%) whereas CLK1(Δ12)-RFP and CLK1(ΔN)-RFP had no significant effect on SRSF1 speckle occupancy (Fig5B). As expected, GFP-SRSF1 localized to nuclear speckles based on the SC35 marker. Such findings indicate that the GFP reporter protein attached to the SR protein did not impact subnuclear localization phenomena. Overall, these results suggest that the large oligomeric form rather than the monomeric or dimeric form is essential for efficient CLK1 entry into and SRSF1 release from nuclear speckles.

Percent exon inclusion is obtained from IMAGEJ analyses of three separate transfection experiments (n=3). P-values are calculated using a one-way ANOVA test (* is P<.0001, ** is P> .05).

### Splicing of *CLK1* is controlled through extensive CLK1 oligomerization

Having shown that high-order oligomerization induced by the N-terminus alters CLK1 and SRSF1 speckle occupancy, we next explored whether these subnuclear changes could affect CLK1 function in the cell. To address this, we monitored the alternative splicing of the mRNA for the *CLK1* gene as a function of recombinant CLK1 expression. Prior studies have shown that CLK1 down-regulates its own cellular expression levels through the exclusion of exon 4, thereby providing a means for tracking the cellular function of CLK1 [32, 33, 35, 36]. We transfected HeLa cells with CLK1-RFP, CLK1(Δ12)-RFP and CLK1(ΔN)-RFP vectors and confirmed their protein expression levels using Western blotting (SI Fig 3). We then isolated RNA from these cells and, using RT-PCR and probes for exons 2 and 5 in the endogenous *CLK1* gene, found that CLK1-RFP strongly induced exon 4 exclusion compared to CLK1(ΔN)-RFP expression (Fig5C). The latter is likely due to the inability of CLK1(ΔN)-RFP to efficiently enter speckles and catalyze Ser-Pro phosphorylation for SR protein activation and release. Most importantly, CLK1(Δ12)-RFP also did not induce exon 4 exclusion but instead exhibited inclusion to the same level as CLK1(ΔN)-RFP suggesting that oligomerization may be important for splicing function. These findings indicate that although the first 100 residues are not essential for generating the Hyper-P state of SRSF1 in our *in vitro* kinetic assays, they serve an important function in trafficking the SR protein in the nucleus for splicing activity. Overall, these results demonstrate that N-terminal sequences supporting oligomerization control CLK1 entry into and SRSF1 release from speckles, a subnuclear transition that plays a role in the autoregulation of CLK1 through splicing of a key alternative exon.

## Discussion

Since the human proteome likely contains many thousands of unique proteins, the phosphorylation of a specific residue within a sea of viable targets by one protein kinase reflects a substantial recognition problem. Early studies on protein kinase A (PKA) suggested that precision targeting may be limited owing to the size of the active site and the available residues for binding [37, 38]. PKA binds the R-R-X-S sequence motif using several negatively charged amino acids but peptides based on this “consensus sequence” that are expected to represent the physiological protein substrate bind with relatively weak affinities [39–41]. To provide a higher layer of selectivity, protein kinases have been found to recruit docking regions outside the active site (within the kinase domain) or within contiguous domains or protein subunits that recognize additional residues up- or down-stream of the consensus sequence [42, 43]. Furthermore, many protein kinases are also localized near their substrates through adapter proteins that also position them among other enzymes in dedicated signaling cascades [44, 45]. For the majority of protein kinases, folded domains are typically used to facilitate these critical protein-protein interactions. However, the CLKs part company with these traditional paradigms by utilizing intrinsically disordered rather than folded domains to recognize their protein partners. In this new study, we provide insights into how CLK1 uses its disordered, non-conserved N-terminus not only to recognize its physiological substrate SRSF1 with high affinity but also to target vital substructures in the nucleus for SR protein-dependent splicing regulation.

Although necessary for recognizing the SR protein family of substrates, how the non-conserved, unstructured N-terminus in CLK1 introduces specificity for splicing function is not well understood. We found in prior studies that CLK1 lacking the full N-terminus (CLK1(ΔN)) interacts with very poor affinity to the traditional SR protein SRSF1. While the K_m_ for SRSF1 is very low (100 nM), indicative of a good protein substrate, removal of the N-terminus results in a non-saturable, steady-state kinetic profile consistent with a K_m_ well in excess of the soluble concentration achievable for the substrate (K_m_ > 2000 nM). Furthermore, although CLK1(ΔN) is a folded domain displaying catalytic activity toward Arg-Ser dipeptides [11, 28], it is not capable of phosphorylating Ser-Pro dipeptides in SRSF1, a critical modification for splicing function. Thus, the N-terminus provides the necessary platform to both recognize SRSF1 and activate the neighboring kinase domain. We now show that residues directly flanking the kinase domain are sufficient to impart essential Ser-Pro dipeptide phosphorylation activity. We suspect that the induction of this capability in the kinase domain could be due to two possible phenomena. First, residues in block 3 of the N-terminus could bind directly to the kinase domain (N-K interactions) inducing a form that better engages Ser-Pro dipeptides from the RS domain in the CLK1 active site (Fig6). We found that these dipeptides are modified later in the reaction time course and, thus, may require greater activation than the Arg-Ser dipeptides, an enhancement provided by a portion of the N-terminus. Second, residues in block 3 are likely to interact directly with the RS domain in SRSF1 and, thus, may position the Ser-Pro dipeptides for optimal attack by the nearby active site of the CLK1 kinase domain. Addition of block 3 not only endows the kinase domain with the ability to hyper-phosphorylate SRSF1 but also provides an interaction surface for the SR protein as evidenced by an improved steady-state kinetic profile for CLK1(Δ12) compared to CLK1(ΔN). Finally, it is also possible that both modes of activation through N-K interactions and RS domain positioning could lead to initiation of the basal hyper-phosphorylation activity and SRSF1 recognition.

An important discovery from our analyses is that high-order CLK1 oligomerization is not necessary for inducing Ser-Pro hyper-phosphorylation of SRSF1. We found that a large excision of N-terminal amino acids reduces CLK1 from a large oligomer, with perhaps 20 or more subunits (M_r_ > 1000 kDa), to a small dimer or possibly an elongated monomer without abolishing generation of the Hyper-P state of SRSF1. However, we found that these oligomerization sequences in the N-terminus are important for CLK1 entry into nuclear speckles (Fig6). Once in speckles, CLK1 can then modify SRSF1 on Ser-Pro dipeptides in the RS domain, a transformation that liberates the SR protein for engagement with the spliceosome. We showed that the oligomerization sequences, while not necessary for attainment of the Hyper-P state of SRSF1, induces the alternative splicing of exon 4 in CLK1, thereby acting to control the cellular protein levels of the kinase. How CLK1 gains entry into speckles to control this regulatory loop is not fully understood. SR proteins are thought to localize in these non-membrane compartments owing to their RS domains that induce large particles suitable to the aggregated environment of speckles. Since the N-termini of CLKs contain numerous disorder-promoting residues similar to the RS domains in SR proteins (Fig1A), it is possible that the forces that increase SR protein aggregation are also at play with the CLKs. Furthermore, it is possible that CLK1 gains access to speckles by using oligomerization sequences to further stabilize interactions with SR proteins that possess speckle-targeting residues. Thus, SR proteins may act to sequester CLKs in speckles for RS domain phosphorylation. Regardless of any specific mechanism, the data presented here suggest that the CLK1 N-terminus uses its oligomerization sequences as a “speckle-recruiting module” to enter a large pool of SR proteins, possibly by mimicking properties of these splicing factors or “piggybacking” with SR proteins, and then subsequently using a separate N-terminal segment to activate the kinase domain for CLK1-specific modifications in SRSF1 (Fig6). It will be interesting in future studies to see how other CLKs utilize their unique N-termini to facilitate subnuclear targeting for SR protein activation and splicing function.

## Conclusion

In this study, we showed that CLK1 uses both structured and unstructured components to enter membrane-less, nuclear substructures known as speckles and catalyze multi-site phosphorylation of SR proteins. While the kinase domain in CLK1 possesses the necessary catalytic machinery to support basal phosphorylation, its activity is considerably weak and requires an intrinsically disordered N-terminus to activate the domain possibly by inducing a productive conformation of the RS domain of the SR protein. We showed herein that the N-terminus can be divided into separate regions that regulate both the core phosphorylation activity and quaternary structure of CLK1. Although high-order oligomerization is not required for activating the unique phosphorylation mechanism encoded in the kinase domain, it is necessary for the nuclear trafficking of CLK1 for SR protein targeting. Disconnecting this oligomerization feature did not abrogate phosphorylation *in vitro* suggesting that residues directly flanking the kinase domain are sufficient for catalytic activation but instead significantly impaired the auto-regulatory splicing function of CLK1. These findings demonstrate that CLKs partition non-conserved residues in a large, disordered N-terminal region not only to activate an otherwise repressed kinase domain but also to recruit it to specific areas of the nucleus for SR protein processing and splicing function.

## Acknowledgements

The authors thank Kendra Hailey for advice, discussion, and technical help. This work was supported by grant GM067969 from the National Institute of Health.

## Supporting information

**SI Fig 1. SDS-PAGE analyses of selected SEC fractions for CLK1 and its deletion constructs and SEC of GST-N(Δ3) in the presence of TCEP.** A) Selected major peak fractions from the S200 column displayed in Fig1F for CLK1, CLK1(Δ1), CLK1(Δ12) and CLK1(ΔN) run on SDS-PAGE. Fractions from oligomer peaks for CLK1 and CLK1(Δ1) and monomer/dimer peaks for CLK1(Δ12) and CLK1(ΔN) are displayed. B) Selected peak fractions from the S200 column displayed in Fig2E for GST-N(Δ3) run on SDS-PAGE. C) Treatment of GST-N(Δ3) with reducing agents. DLS of GST-N(Δ3) is recorded in the absence and presence of incubation with 50 mM TCEP.

**SI Fig 2. CLK1 phosphorylates three Ser-Pro dipeptides in the RS domain of SRSF1.** The sequence of the C-terminal RS domain in SRSF1 and the critical Ser-Pro dipeptides phosphorylated by CLK1 are displayed SRPK1 largely phosphorylates only Arg-Ser dipeptides in the N-terminal portion of the RS domain.

**SI Fig 3. Expression of CLK1-RFP, CLK1(Δ12)-RFP and CLK1(ΔN)-RFP in HeLa cells.** Plasmids encoding CLK1-RFP, CLK1(Δ12)-RFP and CLK1(ΔN)-RFP were transfected into HeLa cells and protein expression levels were assessed using anti-FLAG antibodies.

## References

1. Colwill, K., Pawson, T., Andrews, B., Prasad, J., Manley, J. L., Bell, J. C. & Duncan, P. I. (1996) The Clk/Sty protein kinase phosphorylates SR splicing factors and regulates their intranuclear distribution, EMBO J. 15, 265–75.

2. Colwill, K., Feng, L. L., Yeakley, J. M., Gish, G. D., Caceres, J. F., Pawson, T. & Fu, X. D. (1996) SRPK1 and Clk/Sty protein kinases show distinct substrate specificities for serine/arginine-rich splicing factors, J Biol Chem. 271, 24569–75.

3. Stojdl, D. F. & Bell, J. C. (1999) SR protein kinases: the splice of life, Biochem Cell Biol. 77, 293–8.

4. Zhou, Z. & Fu, X. D. (2013) Regulation of splicing by SR proteins and SR protein-specific kinases, Chromosoma. 122, 191–207.

5. Cho, S., Hoang, A., Sinha, R., Zhong, X. Y., Fu, X. D., Krainer, A. R. & Ghosh, G. (2011) Interaction between the RNA binding domains of Ser-Arg splicing factor 1 and U1-70K snRNP protein determines early spliceosome assembly, Proc Natl Acad Sci U S A. 108, 8233–8.

6. Keshwani, M. M., Aubol, B. E., Fattet, L., Ma, C. T., Qiu, J., Jennings, P. A., Fu, X. D. & Adams, J. A. (2015) Conserved proline-directed phosphorylation regulates SR protein conformation and splicing function, Biochem J. 466, 311–22.

7. Zahler, A. M., Lane, W. S., Stolk, J. A. & Roth, M. B. (1992) SR proteins: a conserved family of pre-mRNA splicing factors, Genes Dev. 6, 837–47.

8. Ge, H. & Manley, J. L. (1990) A protein factor, ASF, controls cell-specific alternative splicing of SV40 early pre-mRNA in vitro, Cell. 62, 25–34.

9. Krainer, A. R., Conway, G. C. & Kozak, D. (1990) The essential pre-mRNA splicing factor SF2 influences 5’ splice site selection by activating proximal sites, Cell. 62, 35–42.

10. Fu, X. D. & Maniatis, T. (1992) The 35-kDa mammalian splicing factor SC35 mediates specific interactions between U1 and U2 small nuclear ribonucleoprotein particles at the 3’ splice site, Proc Natl Acad Sci U S A. 89, 1725–9.

11. Aubol, B. E., Plocinik, R. M., Keshwani, M. M., McGlone, M. L., Hagopian, J. C., Ghosh, G., Fu, X. D. & Adams, J. A. (2014) N-terminus of the protein kinase CLK1 induces SR protein hyperphosphorylation, Biochem J. 462, 143–52.

12. Aubol, B. E., Plocinik, R. M., Hagopian, J. C., Ma, C. T., McGlone, M. L., Bandyopadhyay, R., Fu, X. D. & Adams, J. A. (2013) Partitioning RS domain phosphorylation in an SR protein through the CLK and SRPK protein kinases, J Mol Biol. 425, 2894–909.

13. Muraki, M., Ohkawara, B., Hosoya, T., Onogi, H., Koizumi, J., Koizumi, T., Sumi, K., Yomoda, J., Murray, M. V., Kimura, H., Furuichi, K., Shibuya, H., Krainer, A. R., Suzuki, M. & Hagiwara, M. (2004) Manipulation of alternative splicing by a newly developed inhibitor of Clks, J Biol Chem. 279, 24246–54.

14. Ngo, J. C., Chakrabarti, S., Ding, J. H., Velazquez-Dones, A., Nolen, B., Aubol, B. E., Adams, J. A., Fu, X. D. & Ghosh, G. (2005) Interplay between SRPK and Clk/Sty kinases in phosphorylation of the splicing factor ASF/SF2 is regulated by a docking motif in ASF/SF2, Mol Cell. 20, 77–89.

15. Aubol, B. E., Keshwani, M. M., Fattet, L. & Adams, J. A. (2018) Mobilization of a splicing factor through a nuclear kinase-kinase complex, Biochem J. 475, 677–690.

16. Lindberg, M. F. & Meijer, L. (2021) Dual-Specificity, Tyrosine Phosphorylation-Regulated Kinases (DYRKs) and cdc2-Like Kinases (CLKs) in Human Disease, an Overview, Int J Mol Sci. 22.

17. Naro, C., Bielli, P. & Sette, C. (2021) Oncogenic dysregulation of pre-mRNA processing by protein kinases: challenges and therapeutic opportunities, The FEBS journal. 288, 6250–6272.

18. Martín Moyano, P., Němec, V. & Paruch, K. (2020) Cdc-Like Kinases (CLKs): Biology, Chemical Probes, and Therapeutic Potential, Int J Mol Sci. 21.

19. George, A., Aubol, B. E., Fattet, L. & Adams, J. A. (2019) Disordered protein interactions for an ordered cellular transition: Cdc2-like kinase 1 is transported to the nucleus via its Ser-Arg protein substrate, J Biol Chem. 294, 9631–9641.

20. Lai, M. C., Lin, R. I. & Tarn, W. Y. (2001) Transportin-SR2 mediates nuclear import of phosphorylated SR proteins, Proc Natl Acad Sci U S A. 98, 10154–9.

21. Maertens, G. N., Cook, N. J., Wang, W., Hare, S., Gupta, S. S., Oztop, I., Lee, K., Pye, V. E., Cosnefroy, O., Snijders, A. P., KewalRamani, V. N., Fassati, A., Engelman, A. & Cherepanov, P. (2014) Structural basis for nuclear import of splicing factors by human Transportin 3, Proc Natl Acad Sci U S A. 111, 2728–33.

22. Aubol, B. E., Serrano, P., Fattet, L., Wuthrich, K. & Adams, J. A. (2018) Molecular interactions connecting the function of the serine-arginine-rich protein SRSF1 to protein phosphatase 1, J Biol Chem. 293, 16751–16760.

23. Aubol, B. E., Hailey, K. L., Fattet, L., Jennings, P. A. & Adams, J. A. (2017) Redirecting SR Protein Nuclear Trafficking through an Allosteric Platform, J Mol Biol. 429, 2178–2191.

24. Aubol, B. E., Fattet, L. & Adams, J. A. (2020) A conserved sequence motif bridges two protein kinases for enhanced phosphorylation and nuclear function of a splicing factor, *FEBS J*, May 2. doi: 10.1111/febs.15351.

25. Aubol, B. E., Wu, G., Keshwani, M. M., Movassat, M., Fattet, L., Hertel, K. J., Fu, X. D. & Adams, J. A. (2016) Release of SR Proteins from CLK1 by SRPK1: A Symbiotic Kinase System for Phosphorylation Control of Pre-mRNA Splicing, Mol Cell. 63, 218–228.

26. Aubol, B. E., Fattet, L. & Adams, J. A. (2021) A conserved sequence motif bridges two protein kinases for enhanced phosphorylation and nuclear function of a splicing factor, FEBS J. 288, 566–581.

27. Keshwani, M. M., Hailey, K. L., Aubol, B. E., Fattet, L., McGlone, M. L., Jennings, P. A. & Adams, J. A. (2015) Nuclear protein kinase CLK1 uses a non-traditional docking mechanism to select physiological substrates, Biochem J. 472, 329–38.

28. Bullock, A. N., Das, S., Debreczeni, J. E., Rellos, P., Fedorov, O., Niesen, F. H., Guo, K., Papagrigoriou, E., Amos, A. L., Cho, S., Turk, B. E., Ghosh, G. & Knapp, S. (2009) Kinase domain insertions define distinct roles of CLK kinases in SR protein phosphorylation, Structure. 17, 352–62.

29. Duncan, P. I., Howell, B. W., Marius, R. M., Drmanic, S., Douville, E. M. & Bell, J. C. (1995) Alternative splicing of STY, a nuclear dual specificity kinase, J Biol Chem. 270, 21524–31.

30. Aubol, B. E. & Adams, J. A. (2011) Applying the brakes to multisite SR protein phosphorylation: substrate-induced effects on the splicing kinase SRPK1, Biochemistry. 50, 6888–900.

31. Zhou, J. & Adams, J. A. (1997) Participation of ADP dissociation in the rate-determining step in cAMP-dependent protein kinase, Biochemistry. 36, 15733–8.

32. Duncan, P. I., Stojdl, D. F., Marius, R. M. & Bell, J. C. (1997) In vivo regulation of alternative pre-mRNA splicing by the Clk1 protein kinase, Mol Cell Biol. 17, 5996–6001.

33. Ninomiya, K., Kataoka, N. & Hagiwara, M. (2011) Stress-responsive maturation of Clk1/4 pre-mRNAs promotes phosphorylation of SR splicing factor, J Cell Biol. 195, 27–40.

34. Cha, S. (1970) Kinetic behavior at high enzyme concentrations. Magnitude of errors of Michelis-Menten and other approximations, J Biol Chem. 245, 4814–8.

35. Uzor, S., Zorzou, P., Bowler, E., Porazinski, S., Wilson, I. & Ladomery, M. (2018) Autoregulation of the human splice factor kinase CLK1 through exon skipping and intron retention, Gene. 670, 46–54.

36. Sakuma, M., Iida, K. & Hagiwara, M. (2015) Deciphering targeting rules of splicing modulator compounds: case of TG003, BMC Mol Biol. 16, 16.

37. Knighton, D. R., Zheng, J. H., Ten Eyck, L. F., Ashford, V. A., Xuong, N. H., Taylor, S. S. & Sowadski, J. M. (1991) Crystal structure of the catalytic subunit of cyclic adenosine monophosphate-dependent protein kinase [see comments], Science. 253, 407–14.

38. Knighton, D. R., Xuong, N. H., Taylor, S. S. & Sowadski, J. M. (1991) Crystallization studies of cAMP-dependent protein kinase. Cocrystals of the catalytic subunit with a 20 amino acid residue peptide inhibitor and MgATP diffract to 3.0 A resolution, J Mol Biol. 220, 217–20.

39. Kemp, B. E., Benjamini, E. & Krebs, E. G. (1976) Synthetic hexapeptide substrates and inhibitors of 3’:5’-cyclic AMP-dependent protein kinase, Proc Natl Acad Sci U S A. 73, 1038–42.

40. Kemp, B. E., Graves, D. J., Benjamini, E. & Krebs, E. G. (1977) Role of multiple basic residues in determining the substrate specificity of cyclic AMP-dependent protein kinase, J Biol Chem. 252, 4888–94.

41. Adams, J. A. & Taylor, S. S. (1992) Energetic limits of phosphotransfer in the catalytic subunit of cAMP-dependent protein kinase as measured by viscosity experiments, Biochemistry. 31, 8516–22.

42. Roskoski, R., Jr. (2012) ERK1/2 MAP kinases: structure, function, and regulation, Pharmacol Res. 66, 105–43.

43. Remenyi, A., Good, M. C. & Lim, W. A. (2006) Docking interactions in protein kinase and phosphatase networks, Curr Opin Struct Biol. 16, 676–85.

44. Pawson, T. & Scott, J. D. (1997) Signaling Through Scaffold, Anchoring, and Adaptor Proteins, Science. 278, 2075–2080.

45. Kondo, Y., Paul, J. W., 3rd, Subramaniam, S. & Kuriyan, J. (2021) New insights into Raf regulation from structural analyses, Curr Opin Struct Biol. 71, 223–231.

